# A novel *de novo* FEM1C variant is linked to neurodevelopmental disorder with absent speech, pyramidal signs, and limb ataxia

**DOI:** 10.1101/2022.04.24.489208

**Authors:** Abhishek Anil Dubey, Magdalena Krygier, Natalia A. Szulc, Karolina Rutkowska, Joanna Kosińska, Agnieszka Pollak, Małgorzata Rydzanicz, Tomasz Kmieć, Maria Mazurkiewicz-Bełdzińska, Wojciech Pokrzywa, Rafał Płoski

## Abstract

The principal component of the protein homeostasis network is the ubiquitin-proteasome system. Ubiquitination is mediated by an enzymatic cascade involving, i.e., E3 ubiquitin ligases, many of which belong to the cullin-RING ligases family. Genetic defects in the ubiquitin-proteasome system components, including cullin-RING ligases, are known causes of neurodevelopmental disorders. Using exome sequencing to diagnose a pediatric patient with developmental delay, pyramidal signs, and limb ataxia, we identified a *de novo* missense variant c.376G>C; p.(Asp126His) in the *FEM1C* gene encoding a cullin-RING ligase substrate receptor. This variant alters a conserved amino acid located within a highly constrained coding region and is predicted as pathogenic by most *in silico* tools. In addition, a *de novo FEM1C* mutation of the same residue p.(Asp126Val) was associated with an undiagnosed developmental disorder, and the relevant variant (FEM1C^Asp126Ala^) was found to be functionally compromised *in vitro*. Our computational analysis showed that FEM1C^Asp126His^ hampers protein substrate binding. To further assess its pathogenicity, we used the nematode *Caenorhabditis elegans*. We found that the FEM-1^Asp133His^ animals (expressing variant homologous to the *FEM1C* p.(Asp126Val)) had normal muscle architecture yet impaired mobility. Mutant worms were sensitive to the acetylcholinesterase inhibitor aldicarb but not levamisole (acetylcholine receptor agonist), showing that their disabled locomotion is caused by synaptic abnormalities and not muscle dysfunction. In conclusion, we provide the first evidence from an animal model suggesting that a mutation in the evolutionarily conserved FEM1C Asp126 position causes a neurodevelopmental disorder in humans.

## Introduction

Neurodevelopmental disorders (NDDs) are a group of neurological and psychiatric conditions that arise from disruption of brain development. The defining clinical features are impairments of motor skills, cognition, communication, and/or behavior. NDDs encompass a wide range of disabilities, such as developmental delay (DD), intellectual disability (ID), cerebral palsy (CP), autism spectrum disorder (ASD), attention deficit hyperactivity disorder (ADHD), epilepsy, and others. The etiology of NDDs is heterogeneous and includes multifactorial, acquired, and monogenic causes, the latter often caused by damaging *de novo* mutations (Deciphering Developmental Disorders Study, 2017; Thapar et al., 2017). Interestingly, whereas the pathogenesis of brain dysfunction in NDDs is still poorly understood, at least 62 forms of NDDs are caused by mutations in genes encoding components of the ubiquitin-proteasome system (UPS), especially E3 ubiquitin ligases (Ebstein et al., 2021).

The UPS orchestrates protein degradation in a process known as ubiquitination, where a small protein ubiquitin is covalently attached to its target (Kerscher et al., 2006). Ubiquitination is mediated by an enzymatic cascade involving ubiquitin-activating (E1), ubiquitin-conjugating (E2), and ubiquitin ligase (E3) enzymes (Komander, 2009). The proteasome complex recognizes ubiquitinated proteins and, through proteolysis, degrades them into short peptides that can be further processed (Hochstrasser, 1996). The majority of E3s belong to the RING family, and within it, the largest subfamily is the cullin-RING ligases (CRLs) (Nguyen et al., 2017). CRL complexes are large molecular machineries consisting of a cullin protein that acts as a scaffold, a RING (Really Interesting New Gene) box protein coordinating ubiquitination, a substrate receptor that recruits the target protein, and adaptor proteins that link the substrate receptor to the cullin (Nguyen et al., 2017).

FEM1A, FEM1B, and FEM1C are evolutionarily conserved substrate recognition subunits of cullin 2-RING E3, which selectively bind their target proteins via a specific sequence motif called a degron (Dankert et al., 2017) (Fig. 1A). Degron is a destabilizing short linear motif that may be located anywhere in the protein sequence, including its N-/C-terminus which may contain motifs belonging to the N-/C-degron pathways, respectively (Varshavsky, 2019). Recent studies demonstrated that FEM1C receptors recognize various arginine-terminated motifs (Arg/C-degrons) (Koren et al., 2018; Lin et al., 2018), directing them to proteasomal degradation. The known FEM1C substrates include SIL1, a nucleotide exchange factor for the BiP endoplasmic reticulum chaperone associated with Marinesco-Sjögren syndrome - an autosomal recessive cerebellar ataxia (Anttonen et al., 2005; Senderek et al., 2005).

**Figure 1.**
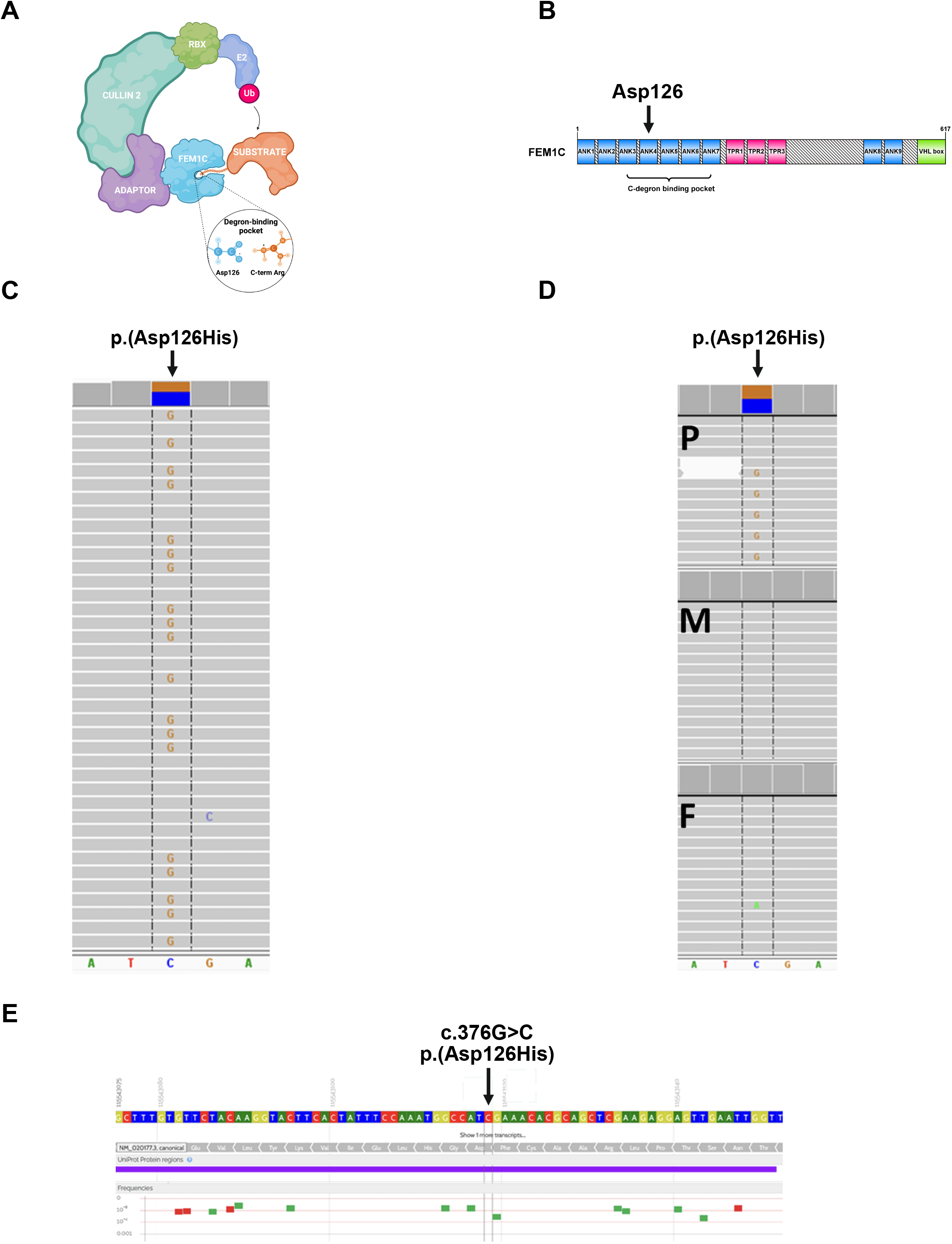
Results of the genetic study. **A. Components of CRL complex on the example of FEM1C. B**. Scheme of the domain architecture of human FEM1C visualized using the IBS software (Liu et al., 2015). ANK - ankyrin repeat; TPR - tetratricopeptide repeat; VHL box - von Hippel– Lindau box. Modified from Chen et al. (2021). **C**. Missense p.(Asp126His) variant in *FEM1C* gene identified in the proband by WES (coverage 119x, reads with alternate allele 57). **D**. Amplicon deep sequencing-based family study of p.(Asp126His) (P - proband: coverage 6459x, reads with alternate allele 3097, M - mother: coverage 6171x, reads with alternate allele 0, F – father: coverage 5931, reads with alternate allele 0). **E**. Overview of genetic variation in the proximity of p.(Asp126His) in a healthy population (GnomAD and Bravo cohorts, based on https://varsome.com).

We describe a nine-year-old boy with severe DD, lack of speech, pyramidal signs, and limb ataxia. Whole exome sequencing (WES) identified a *de novo* missense variant c.376G>C; p.(Asp126His) in the *FEM1C* gene. *De novo* mutation in the same residue, FEM1C^Asp126Val^, was previously associated with an uncharacterized DD (Deciphering Developmental Disorders Study, 2017), pointing to the importance of this position in FEM1C function in humans. Yet another such variant (FEM1C Asp126Ala) was shown to be functionally relevant by *in vitro* studies (Chen et al., 2021; Yan et al., 2021). Furthermore, a *de novo* mutation at the corresponding amino acid residue in another member of the FEM1 family, FEM1B^Arg126Gln^ (closely related to FEM1C), was associated with syndromic intellectual disability (Lecoquierre et al., 2019). The FEM1C disease-linked variant Asp126His occurs within its degron-binding pocket and impacts a residue directly involved in binding the critical C-terminus arginine of FEM1C substrates (Chen et al., 2021; Yan et al., 2021) (Fig. 1A, B).

Here we show that the pathogenicity of the p.(Asp126His) variant found in the patient is supported by studies of nematode *Caenorhabditis elegans* with a corresponding mutation in FEM-1 (a worm homolog of human FEM1C). Our results provide the first evidence of an NDD associated with a novel *FEM1C* mutation that has functional consequences for the nervous system in the *C. elegans* model.

## Results

### Exome sequencing and family studies

ES performed in the proband revealed three ultra-rare (0 frequency in gnomAD v3.1.2 https://gnomad.broadinstitute.org and an in-house database of >5500 Polish exomes) heterozygous missense variants in *CSMD2, RALYL* and *FEM1C* genes which were selected for family studies. Variants in both *CSMD2* and *RALYL* were found to be inherited from a healthy mother and father and thus were considered as not disease-causing. The *FEM1C* variant (hg38, chr5:115543118-C>G, NM_020177.3:c.376G>C, p.(Asp126His); Fig. 1C) was absent in both parents, indicating a *de novo* event (Fig. 1D). The *FEM1C* p.(Asp126His) was predicted as damaging by 13 different pathogenicity predictors implemented in the Varsome web server (https://varsome.com), including BayesDel addAF, BayesDel noAF, EIGEN, EIGEN PC, FATHMM-MKL, FATHMM-XF, LRT, M-CAP, MutPred, MutationTaster, PROVEAN, PrimateAI, SIFT, and as “Tolerated” by six other predictors (DEOGEN2, FATHMM, LIST-S2, MVP, Mutation assessor, SIFT4G). The c.376G position is highly conserved (PhastCons100way score 1.0, GERP++ value 5.5), and the p.(Asp126His) variant affects the conserved Asp126 residue located in the ankyrin repeat domain ANK4 (Fig. 1B) (phyloP100way = 7.9). Moreover, *FEM1C* p.(Asp126His) variant is located in the highly constrained coding region (CCR=98.6). The scarcity of coding variants in the proximity of p.(Asp126His), characteristic of CCR, is illustrated in Fig. 1E, based on the gnomAD and Bravo datasets.

### FEM1C^Asp126His^ disrupts degron binding

Recent structural studies on FEM1C revealed a crucial role for Asp126 in degron recognition and its direct interactions with the Arg residue of the Arg/C-degron substrates. Isothermal titration calorimetric performed by Yan and colleagues showed that Asp126Ala mutation in FEM1C lowers its binding affinity towards Arg/C-degron substrates ∼2-fold (Yan et al., 2021). In addition, the Global Protein Stability assay conducted by Chen and colleagues revealed that FEM1C double mutant Asp77Ala/Asp126Ala failed to induce the degradation of three different Arg/C-degron substrates (Chen et al., 2021), further indicating the importance of Asp126 in FEM1C functioning. Therefore, we hypothesized that Asp126His mutation carried by the patient might also abrogate the degron binding by its repulsive interactions with C-terminal arginine and thus underlie the disease by impairing ubiquitination and turnover of FEM1C substrates. To gain insights into the binding modes of FEM1C and its mutated variants - Asp126His, Asp126Ala, and Asp126Val (as in another patient with DD), we performed the peptide-protein docking using a novel protocol utilizing the AlphaFold2 modeling (Tsaban et al., 2022). To this end, we docked the SIL1 C-terminus peptide to FEM1C and its single-point mutants, and since the experimental structure of FEM1C bound to the Arg/C-degron of its native substrate SIL1 was resolved (PDB ID: 6LBN) (Chen et al., 2021), we checked whether the obtained AlphaFold2 models managed to recapitulate the interactions present in the experimentally resolved complex. We obtained such similar complexes only for the wild-type FEM1C, where 14/17 (model 1) or 13/17 (model 4’) of native interactions were recapitulated (Fig. 2A), including the interaction of FEM1C Asp126 with the C-terminal arginine of SIL1 (Fig. 2B-C). Remarkably, none of the mutants exhibited the peptide docked to its degron-binding pocket (Fig 2A, Table S1), although the root-mean-square deviation (RMSD), a widely-used metric to assess the similarity between structures, indicated that the degron-binding region of all AlphaFold2 FEM1C models was predicted as similarly folded to the experimental structure (RMSD in ranges of 0.322 – 0.431 Å) (Supp. Table 1). These results confirm the critical role of FEM1C Asp126 residue in the Arg/C-degron recognition and implicate the pathogenic character of both Asp126His and Asp126Val mutations which likely induce severe changes in the environment of the FEM1C degron-binding pocket.

**Figure 2.**
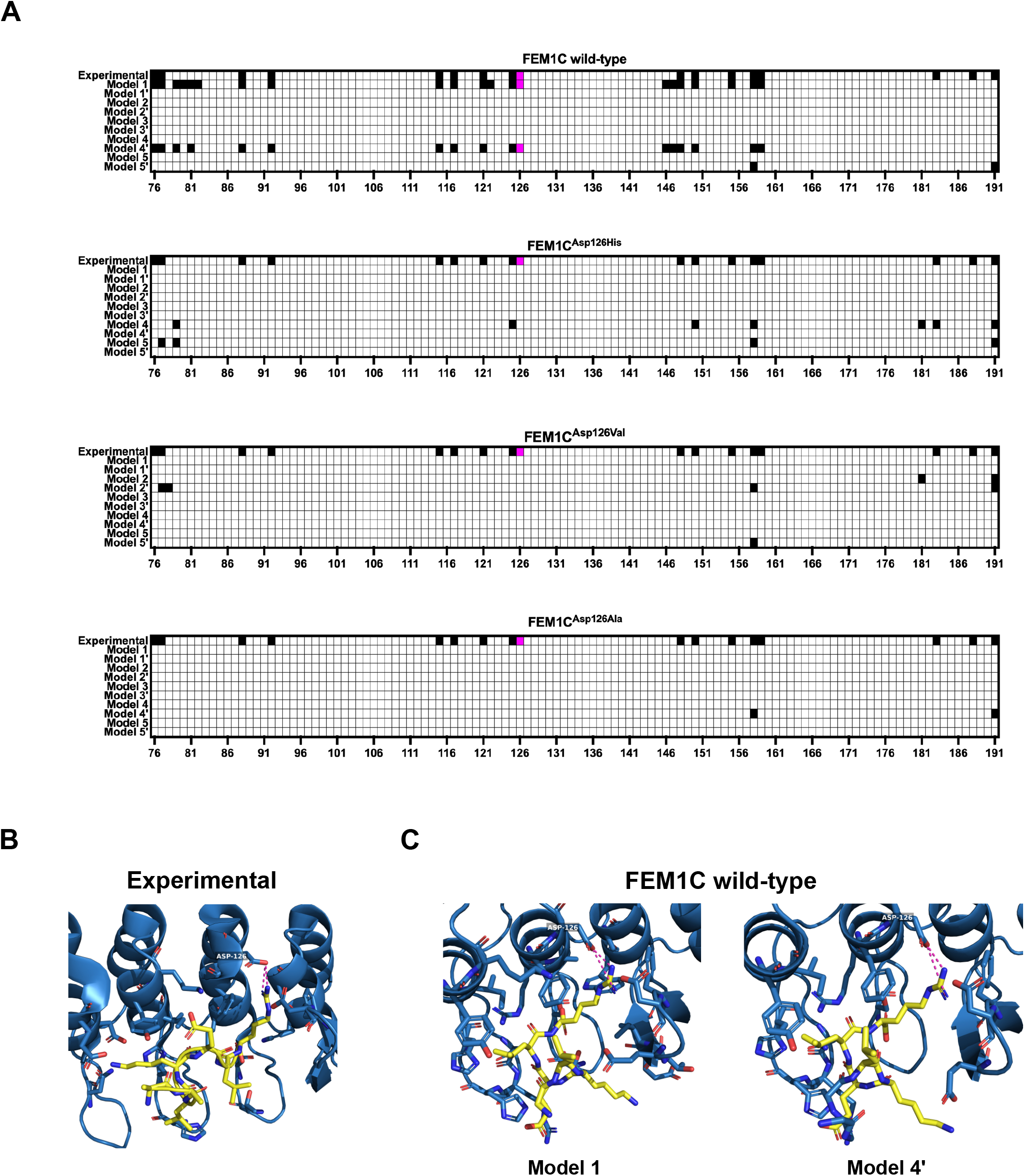
Arg/C-degron from SIL1 docked to human FEM1C and its variants. **A**. Contact heatmaps between FEM1C and its variants and the LKELR Arg/C-degron of SIL1 obtained from the AlphaFold2 protein-peptide docking; experimental – interactions present in the resolved structure of the complex (PDB ID: 6LBN); interaction between Asp126 and the C-terminal arginine of SIL1 is colored in magenta. Contacts were defined as not exceeding the 4 Å distance between any heavy atoms. Data visualized in the GraphPad Prism 9 software. **B**. Visualization of interactions between Arg/C-degron of SIL1 with FEM1C from the experimentally resolved complex; blue - FEM1C, yellow – LKELR Arg/C-degron of SIL1, magenta - hydrogen bonds between FEM1C Asp126 and C-terminal arginine of SIL1. Visualized in the PyMOL software (Schrödinger). **C**. Visualization of interactions between Arg/C-degron of SIL1 with FEM1C from two AlphaFold2 models (of wild-type FEM1C) where the interaction of the C-terminal arginine of SIL1 with Asp126 of FEM1C was recapitulated; blue - FEM1C, yellow – LKELR Arg/C-degron of SIL1, magenta - hydrogen bonds between FEM1C Asp126 and C-terminal arginine of SIL1. Visualization was done in the PyMOL software (Schrödinger).

### The nematode *C. elegans* as a model to study the functional consequences of the FEM1C p.(Asp126His) mutation

The degron-binding pocket of FEM1C is located within its N-terminal ankyrin domain (Chen et al., 2021; Yan et al., 2021) (Fig. 1B), a well-conserved motif in all FEM1 proteins and their orthologs in other organisms, including *C. elegans* nematode. *C. elegans* is a well-established model organism to study multiple human diseases, including neurodegeneration and ataxias (Alexander et al., 2014; Sorkaç et al., 2016). Therefore, we used worms to investigate the physiological impact of the FEM1C p.(Asp126His) mutation. The sequence alignment of degron-binding regions of FEM1C and its *C. elegans* ortholog FEM-1 showed high evolutionary conservation, including the critical Asp126 position (Asp133 in worms) (Fig. 3A; left panel). We also compared the structures of both proteins, as protein folding directly underlies molecular interactions and may point to the conserved functional role. Superposition of the experimental structure of the FEM1C degron-binding region (residues 1-390) and model of its FEM-1 analog obtained from the AlphaFold database (Jumper et al., 2021; Varadi et al., 2021) showed very high structural similarity, evidenced in particular by a low RMSD value of 1.213 Å).

**Figure 3.**
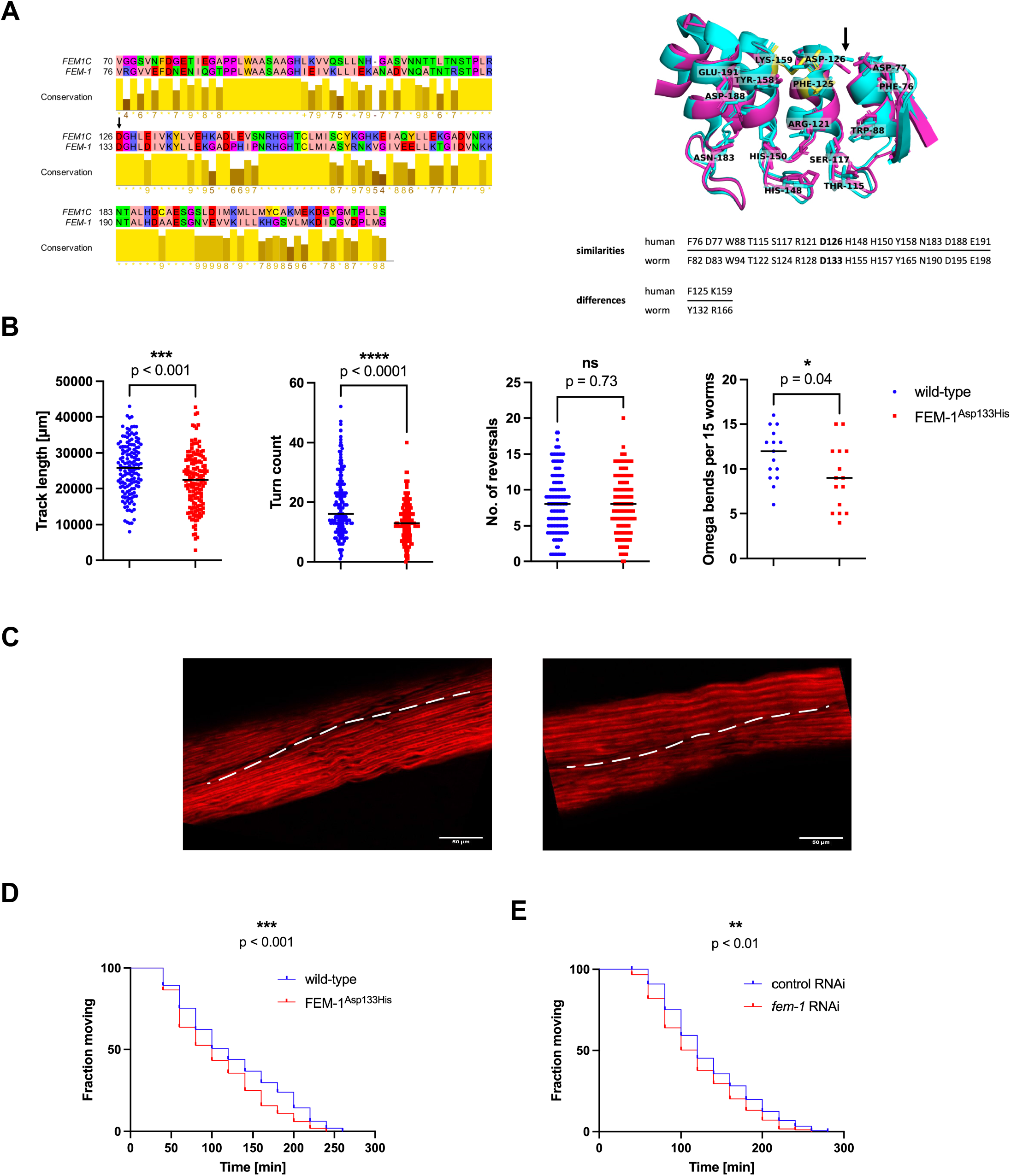
Computational and functional analysis of the *C. elegans fem-1* mutant. **A. Left:** Global alignment of binding regions of human FEM1C (residues 70-220; UniProt ID: <u>Q96JP0</u>) and *C. elegans* FEM-1 (residues 76-227; UniProt ID: <u>P17221</u>). Sequences were aligned using the EMBOSS Needle web server (Madeira et al., 2022) with default parameters and visualized in the Jalview Desktop software (Waterhouse et al., 2009) with residues colored by their physicochemical properties; the arrow indicated Asp126/Asp133 position; **Right:** Comparison of residues involved in SIL1 Arg/C-degron binding in FEM1C (cyan; PDB code: 6LBN) and FEM-1 (magenta; AlphaFold2 model corresponding to P17221). FEM-1 residues that differ from FEM1C were marked in yellow; the arrow indicated the Asp126/Asp133 position. **B**. Track lengths, turn counts, a number of reversals, and omega bends in wild-type (*n*=146) and FEM-1^Asp133His^ (*n=*148) worms; *N*=3. *n* represents the total number of worms; *N* represents the number of experimental repeats. The stars denote the significance levels per the *p-value* obtained by the Mann–Whitney test. *p-values* are shown adjacent to the respective graphs (ns - not significant, \**p* ≤ 0.05, \*\*\**p* ≤ 0.001, \*\*\*\**p* ≤ 0.0001), dashes represent median of collective data from three independent biological replicates. Data was plotted and analyzed in the GraphPad Prism 9 software. **C**. Phalloidin staining of F-actin-containing I-bands of wild-type and FEM-1^Asp133His^ worms. Dashed lines separate two body wall muscle cells. The scale bar is 50 μm. **D**. Aldicarb sensitivity assay of wild-type (*n*=207) and FEM-1^Asp133His^ (*n*=215) worms; *N*=3. *n* represents the total number of worms; *N* represents the number of experimental repeats. *p-value* was calculated using the Log-rank (Mantel-Cox) test and is shown adjacent to the graph (\*\*\**p* ≤ 0.001). Data was plotted and analyzed in the GraphPad Prism 9 software. **E**. Aldicarb sensitivity assay in neuronal-specific RNAi of *fem-1*, [*fem-1* RNAi (*n*=183)] and vector control [control RNAi (*n*=177)]; *N=3* in the neurons. *n* represents the total number of worms; *N* represents the number of experimental repeats. *p-value* was calculated using the Log-rank (Mantel-Cox) test and is shown adjacent to the graph (\*\**p* ≤ 0.01). Data was plotted and analyzed in the GraphPad Prism 9 software.

Recent structural studies of the FEM1C basis of degron recognition pointed to the critical role of several residues (Chen et al., 2021; Yan et al., 2021). Therefore, we compared their evolutionary conservation to assess the similarity of the degron-binding pocket between humans and *C. elegans*. Among 15 residues of FEM1C indicated as involved in Arg/C-degron binding of its substrate SIL1 (Chen et al., 2021), 13 were conserved both in sequence and structure in FEM-1 (Fig. 3A; right panel). Notably, the two differing residues showed similar physicochemical properties (phenylalanine/tyrosine and lysine/arginine), further supporting the high conservation of the degron binding pocket and indicating a likely similar substrate pool of FEM1C and FEM-1. Thus, the high similarity of the degron-binding pocket between human FEM1C and its worm ortholog, as well as the conservation of Asp126 residue, are strong premises to employ *C. elegans* as a FEM1C p.(Asp126His) disease model.

### *C. elegans* FEM-1^Asp133His^ mutants develop locomotor defects due to neuronal dysfunction

Ataxias are a heterogeneous group of disorders in which cerebellar dysfunction underlies neurological symptoms such as impaired muscle coordination (Shakkottai & Paulson, 2009). As pyramidal signs and limb ataxia were prominent symptoms in our proband, we examined the locomotion behavior of worms expressing the putative disease-causing FEM-1^Asp133His^ mutant protein (hereafter *fem-1* mutants) generated using the CRISPR/Cas9 gene-editing tool (Supp. Fig. 1A). Locomotion is a complex reflection of the functionality of the *C. elegans* nervous system. It encompasses a range of motor activities, such as omega bends (deep bends usually on the ventral side of the body that change the movement direction) and reversals (transition from crawling forward to crawling backward). We used the WormLab system (MBF Bioscience) to measure animal locomotor parameters in wild-type and *fem-1* worms in detail. We found that the *fem-1* mutants had impaired mobility as assessed by track length and turn counts (Fig. 3B). In particular, the *fem-1* mutants experienced difficulty performing omega bends but showed no defects in reversals (Fig. 3B). This suggests a functional deficit of distinct neurons in *fem-1* mutants, as the execution of reversals and omega bends is encoded by separate neurons (Gray et al., 2005).

Moreover, FEM-1 is essential for sperm production, and its deficiency leads to the sterility of *C. elegans* hermaphrodites (Doniach & Hodgkin, 1984). However, expression of FEM-1^Asp133His^ did not affect fertilization, as the number of unhatched eggs (usually accounting for a few percent of all eggs) laid by the mother within 16 hours was similar to that in the wild-type strain, indicating that fertilization and embryo viability remained unchanged (Supp. Fig. 1B). However, *fem-1* mutants laid a lower number of eggs over 16 hours compared to wild-type worms (Supp. Fig. 1C), which may suggest defects related to the development of vulva muscles or the neuronal circuit (six cholinergic and two serotonergic neurons) controlling their contraction, and thus, egg expulsion (Jose et al., 2007).

The organization of sarcomeres and their components in *C. elegans* body wall muscles is a highly evolutionary conserved feature (Gieseler et al., 2017). To determine whether locomotion and egg-laying defects in *fem-1* mutants occur due to the loss of muscle fiber structure, we performed phalloidin staining of F-actin filaments containing I bands (Shai, S., 2016). We did not find myofilament disorganization or reduced muscle cell size in *fem-1* mutants (Fig. 3C), suggesting that the observed deficits in locomotion are not the result of compromised body wall muscle organization.

To confirm that the mobility phenotype observed in the *fem-1* mutants is due to synaptic defects, we examined the rate of worm paralysis in response to aldicarb and levamisole. Acetylcholine (Ach) is necessary for the transmission of an action potential between neurons, and it exerts its action by binding to nicotinic (nAchR) and muscarinic Ach receptors (Sammi et al., 2022; Spensley et al., 2018). Synaptic acetylcholinesterase (AchE) manages neuromuscular transmission by catabolizing Ach. Aldicarb inhibits AchE and induces spastic muscle paralysis due to the accumulation of Ach in the synaptic gap. (Mahoney et al., 2006). Similar paralysis is also induced by levamisole, a nAchR agonist (Qian et al., 2008; Treinin & Jin, 2021) (Supp. Fig. 1D). However, in the aldicarb assay, a higher percentage of paralyzed worms at a given time indicates a higher general Ach neurotransmission, whereas the levamisole test selectively evaluates nAchR activity (Mahoney et al., 2006; Miller et al., 1996; Sammi et al., 2022). We found that the *fem-1* mutants were significantly (p<0.001) more sensitive to aldicarb but not to levamisole (p=0.34) compared to wild-type worms (Fig. 3D and Supp. Fig. 1E). In addition, neuronal-specific RNA (ribonucleic acid) interference (RNAi) depletion of FEM-1 also sensitized animals to aldicarb treatment (p<0.01, Fig. 3E) suggesting that FEM-1^Asp133His^ acts through the decreasing function of the protein. Our data suggest that inefficient turnover of neuronal substrates associated with the mutation in FEM-1 may lead to increased Ach release, e.g., via impaired calcium sensing, overactivity of the machinery responsible for synaptic vesicle release, or excessive amounts of neuromodulatory peptides (Hu et al., 2011; McEwen et al., 2006; Miller et al., 1996; Treinin & Jin, 2021), resulting in aldicarb sensitivity, and motor and egg-laying defects.

## Discussion

We describe a pediatric case of severe DD, lack of speech, pyramidal signs and limb ataxia associated with a novel *de novo* variant p.(Asp126His) in the *FEM1C* gene.

The variant altered a conserved amino acid position and was predicted to be pathogenic by most *in silico* tools. Moreover, the variant is located within a highly constrained coding region, i.e., a region with a scarcity of coding variation among healthy people likely to be enriched in pathogenic variants for autosomal dominant disorders (Havrilla et al., 2019).

Importantly, another *de novo* variant (Asp126Val) in the same residue of FEM1C was associated with a neurodevelopmental disorder (Deciphering Developmental Disorders Study, 2017, additionally reported in Turner et al., 2019). Moreover, a *de novo* variant (Arg>Gln) at homologous residue 126 in another member of the FEM-1 family, FEM1B, was found in a patient with a similar broad phenotype, including global development delay (Deciphering Developmental Disorders Study, 2017; Lecoquierre et al., 2019).

The FEM1C disease-linked mutation p.(Asp126His) occurs within its degron-binding pocket and impacts a residue directly interacting with FEM1C substrates. Structural studies revealed specific FEM1C preferences for binding proteins with arginine (R) or lysine (K) at the preantepenultimate or antepenultimate positions, which indicates that the degron motifs preferred by FEM1C are C-terminal R/K-X-X-R or R/K-X-R sequences (X stands for any amino acid) (Chen et al., 2021; Yan et al., 2021).

A misregulation of SIL1, which is linked to an autosomal recessive cerebellar ataxia and DD (Anttonen et al., 2005; Senderek et al., 2005), is a possible cause of the diseases associated with FEM1C^Asp126His^ or FEM1C^Asp126Val^ mutations. This is supported by experimental evidence showing that FEM1C recognizes a peptide at the C-terminus of SIL1 and that the FEM1C^Asp126Ala^ mutation significantly reduced this interaction (Chen et al., 2021; Yan et al., 2021).

However, R/K-X-X-R or R/K-X-R motifs occur in 292 human proteins, making them all putative substrates of FEM1C (Supp. Table 2). Furthermore, a small amino acid such as glycine (G) or alanine (A) consecutive to the last arginine in the FEM1C substrate may also be tolerated allowing weaker or transient interactions (Chen et al., 2021; Yan et al., 2021). This could extend the pool of putative FEM1C substrates, as there are 14 and 19 proteins in the human proteome yielding R/K-X-X-R-G or R/K-X-R-G and R/K-X-X-R-A or R/K-X-R-A C-terminus motifs, respectively (Supp. Table 2). Furthermore, it has been shown that cleavage by proteolytic enzymes produces new protein ends that can contain destabilizing residues and act as degron motifs (Varshavsky, 2011), which implies that under certain circumstances, the number of potential FEM1C substrates could be much higher.

Pathogenicity of the p.(Asp126His) variant found in our patient is supported by studies of transgenic *C. elegans*, a well-established model organism to study multiple human diseases, including neurodegeneration and ataxias (Alexander et al., 2014; Sorkaç et al., 2016). We found that the patient’s allele reduced animal mobility by affecting omega bends but not reversals. Since the patient’s allele did not affect muscle organization, the reduced mobility was likely of neuronal origin. As the execution of reversals and omega bends is encoded by separate neurons in the worm, the observed functional deficit is likely discreet (Gray et al., 2005).

Consequently, we analyzed the functionality of the nervous system in *fem-1* worms. Using the acetylcholinesterase inhibitor aldicarb, we observed greater sensitivity of mutant animals, as evidenced by a faster rate of muscle paralysis. This finding suggests that locomotor defects induced by a mutation in the *fem-1* gene occur from neuronal synaptic deficits and excessive ACh release in the synaptic cleft. Interestingly, levamisole did not have a similar effect, suggesting that excessive ACh does not exert its effect through nicotinic Ach receptors in mutant worms.

ACh also has a complex role in controlling *C. elegans* egg-laying behavior (Rand, 2007). In this regard, *fem-1* mutants exhibited reduced egg laying, which may be attributed to cholinergic neuronal abnormalities (neurons capable of producing ACh).

As neuronal-specific RNAi depletion of FEM-1 also sensitized animals to aldicarb treatment, the FEM-1^Asp133His^ likely acts through decreasing protein function. However, since the complete loss of function (LOF) of FEM-1 causes sterility of *C. elegans* (Doniach & Hodgkin, 1984), a feature not observed in our FEM-1^Asp133His^ mutant worms, the decrease due to the variant is likely to be partial. Given that human *FEM1C* does not appear to be a haploinsufficient gene (3 LOF variants in GnomAD, pLI=0.63, https://gnomad.broadinstitute.org), we speculate that in the heterozygous patient p.(Asp126His) decreases the protein function by >50%, perhaps acting as a dominant negative hypomorphic allele. Interestingly, a gain of function mechanism was suggested to underlie the disease association of FEM1B Arg126Gln (Manford et al., 2021).

Our results pointing to the functional relevance of FEM1C Asp126His are entirely consistent with previously reported biochemical studies showing that another variant at the same residue - Asp126Ala - abolished or severely abrogated substrate binding by FEM1C (Chen et al., 2021; Yan et al., 2021).

In conclusion, we have identified an NDD associated with a novel FEM1C mutation that exerts functional consequences on the nervous system in a C. elegans model. Since optimal modulation of synaptic activity is balanced by the differential activity of multiple pathways, further targeted genetic and pharmacological interventions using our *fem-1* worms are needed to explore detailed neuronal mechanisms affected by the FEM1C/FEM-1 variant.

## Materials and Methods

The proband family commercially ordered genetic testing for diagnostic purposes. The patient’s legal guardians have written consent to publish genetic and clinical data.

### Clinical report

The patient is the first child of non-consanguineous parents. The family history was unremarkable. The pregnancy was uncomplicated, with normal screening ultrasounds. He was delivered at term via C-section, and all mensuration at birth was normal. Developmental concerns became apparent to the family at around six months of age due to gross motor delay. Neurologic evaluation at nine months of age showed axial hypotonia and increased muscle tone in the lower limbs with brisk tendon reflexes. Magnetic Resonance Imaging (MRI) of the brain with spectroscopy performed at the age of 11 months showed symmetrical enlargement of the posterior horns of the lateral ventricles. The boy could not sit independently until 2.5 years of age and crawl until three years. Brain MRI at the age of four showed mildly enlarged lateral ventricles comparable to the previous examination and small areas of incomplete myelination at the posterior parts of the lateral ventricles. A recent neurological examination at the age of nine years was notable for global developmental delay, lack of speech, ataxia, and pyramidal signs. Anthropometric measurements were normal: weight 26 kg (25 cc), height 134 cm (25 cc), head circumference 54 cm (50-75cc), BMI - 14,5. Normal male sexual organs were observed, with testicles in the scrotum and a normal penis. Head MRI performed at nine years was normal. The patient was cheerful, reacted to simple commands, and communicated using assistive devices. He was not able to walk independently. He stood with support and crossed his legs when held by the caregiver. There was massive nystagmus in all directions of gaze, left divergent strabismus, cerebellar signs in the upper limbs (ataxia, intention tremor, dysmetria, dysdiadochokinesia), axial hypotonia, pyramidal signs (spasticity with brisk deep tendon reflexes in the upper and lower limbs), and bilateral pes planovalgus. Cognitive assessment performed at the age of nine using Stanford-Binet 5 test showed moderate intellectual disability. Due to the lack of verbal speech in the boy, the study only included the development of cognitive skills in the non-verbal areas. Results of metabolic evaluation, including urine organic acid analysis, tandem mass spectrometry screening, and neurotransmitters in cerebrospinal fluid, were normal. Chromosomal microarray did not show any disease-causing copy number variants.

### Genetic study

Exome sequencing (ES) was conducted for the proband only using DNA purified from the whole blood. According to the manufacturer’s instructions, the library was constructed using SureSelect All Human Exon V7 (Agilent Technologies, Cedar Creek, TX, USA). The library was paired-end sequenced (2×100 bp) on HiSeq 1500 (Illumina, San Diego, CA, USA) to the mean depth > 100x (the min. 10x coverage was 96.7%, and 20x - 91.6%). Reads were aligned to the GRCh38 (hg38) reference genome. Data analysis and variants prioritization was performed as previously described (Rydzanicz et al., 2021). Variants considered causative were validated in the proband and studied in his parents by deep amplicon sequencing performed using Nextera XT Kit (Illumina) and paired-end sequencing as described above for WES.

### Peptide-protein docking

Peptide-protein docking was performed as described by Tsaban et al. (2022). Briefly, the sequence of the first 390 residues of FEM1C (corresponding to the experimentally resolved region of FEM1C; PDB ID: 6LBN) or its single-point mutants (Asp126His, Asp126Val, Asp126Ala), followed by a 30 glycine linker and the LKELR Arg/C-degron sequence (corresponding to the C-terminus of SIL1, also present in the 6LBN structure) served as an input for the local version of AlphaFold2 run for the monomer model. Ten models were generated for each variant (in two separate runs); multiple sequence alignment was constructed based on all genetic databases; max_template_date was set to 2022-07-28. Root-mean-square deviation (RMSD) was calculated for C-alpha atoms between the degron-binding pocket of the predicted models and the experimental structure (residues 76-191) using the PyMOL software (Schrödinger).

### Worm maintenance and strains

Worms were maintained on nematode growth medium (NGM) plates seeded with OP50 *E. coli* bacteria at 20°C unless otherwise stated (Brenner, 1974). Wild-type nematodes were the N2 (Bristol) strain. PHX5163 *fem-1(syb5163)* mutant worms were generated at SunyBiotech (http://www.sunybiotech.com) by CRISPR/Cas9 editing using the sgRNAs sg1-CCA TTA AGA GGT GCA TGT TAC GA and sg2-TTA AGA GGT GCA TGT TAC GAT G. The editing was confirmed by sequencing (Supp. Fig. 1A). Strain PHX5163 was outcrossed 2X to N2 to generate strain WOP497. Neuron-specific silencing was carried out using strain TU3311 [uls60 (unc-119p::YFP + unc-119p::sid-1)] (Calixto et al., 2010).

### RNA interference

RNA interference (RNAi) in *C. elegans* was performed using the standard RNAi feeding method and RNAi clone from the Ahringer *C. elegans* RNAi feeding library (Kamath & Ahringer, 2003). For experiments, we used NGM plates supplemented with 1 mM IPTG and 25 μg/μl carbenicillin seeded with HT115 *E. coli* bacteria expressing double-stranded RNA from L4440 plasmid against the *fem-1* gene. N2 and TU3311 worms were placed on freshly prepared RNAi plates as age-synchronized L1 larvae. Worms fed with *E. coli* HT115(DE3) containing empty L4440 plasmid were used as control.

### Worm mobility assay

Day 1 adult worms were placed onto a single NGM plate, and worm movements were recorded for 2 minutes using the WormLab system (MBF Bioscience), keeping the frame rate, exposure time, and gain set to 7.5 frames per second. Parameters such as the track length, turn count, number of reversals, and reversal distance travelled pattern of individual worms were recorded and analyzed using the WormLab software (MBF Bioscience). The assay consisted of three independent biological replicates. 75 worms were recorded for one biological replicate.

### Phalloidin staining

Sarcomere assembly was monitored by staining F-actin with Phalloidin-Atto 390 (Sigma Aldrich) using the protocol described in Romani and Auwerx (2021). Briefly, synchronized day 1 adults were washed from NGM plates seeded with *E. coli* OP50, and the worm pellet was snap-frozen in liquid nitrogen. The pellet was dried in the SpeecVac concentrator, followed by worm permeabilization using 100% ice-cold acetone for 5 minutes. After acetone removal, the worm pellet was stained using Phalloidin-Atto 390 for 30 minutes in the dark. Stained worms were washed twice, mounted onto a 2% (w/v) agarose pad on a glass slide, and imaged on Zeiss LSM800 inverted confocal microscope.

### Aldicarb and levamisole sensitivity assay

Aldicarb and levamisole sensitivity assay was performed as described in Sorkaç et al. (2016) and Felton and Johnson (2014), respectively. Briefly, 20 adult day 1 worms (also these subjected to RNAi) were placed on NGM plates supplemented with 0.5 mM aldicarb or 0.5mM levamisole, and worm paralysis was assessed by their inability to move within 5 seconds in response to touch in the head and tail region.

Paralysis was assessed every 20 minutes until all worms were utterly paralyzed. The assay consisted of three independent biological replicates.

### Egg-laying assay

Synchronized N2 and *fem-1* mutant worms were grown till day 1 adulthood. The egg-laying assay was carried out by transferring a single day 1 adult animal on NGM plate seeded with *E. coli* OP50 and kept at 20ºC for 16 hours. After 16 hours, the worm was removed from the plate, and the eggs were counted. At least 25 animals were scored in each independent set, and the experiment was repeated three times as independent biological replicates.

## Acknowledgments

We thank Dr. Filip Stefaniak for his advice on molecular docking and sharing computational resources. We acknowledge the Caenorhabditis Genetics Centre (funded by the NIH National Centre for Research Resources, P40 OD010440) for the N2 and TU3311 strains and OP50 *E. coli*.

## Supplementary Figure

**A**. Sequencing chromatogram of CRISPR/Cas9-generated FEM-1^Asp133His^ mutation displaying sequence peaks and base calls. Data was visualized in the SnapGene 6 software (Insightful Science; snapgene.com). **B**. Number of unhatched eggs counted after 16 hours of laying for wild-type (*n*=95) and FEM-1^Asp133His^ (*n=*92) worms; *N*=5. *n* represents the total number of worms; *N* represents the number of experimental repeats. The stars denote the level of significance of the *p-value* obtained by the Mann–Whitney test. *p-value* is shown adjacent to the graph (\*\*\**p* ≤ 0.001). Data was plotted and analyzed in the GraphPad Prism 9 software. **C**. Egg-laying assay of wild-type (*n=95*) and FEM-1^Asp133His^ (*n*=92) worms; *N=5. n* represents the total number of worms; *N* represents the number of experimental repeats. *p-value* was calculated obtained by the Mann–Whitney test and is shown adjacent to the graph (ns - not significant). Data was plotted and analyzed in the GraphPad Prism 9 software. **D**. Illustration of aldicarb’s and levamisole’s mechanism of action. The transmission of impulses between neurons is mediated by the synthesis of Ach (magenta star) at the presynapse and its release into the synaptic cleft, where it binds to Ach receptors (only nAchR are shown). Aldicarb (depicted as a stop sign) inhibits AchE (navy pac-man), resulting in the build-up of Ach, which causes muscle paralysis. Levamisole (yellow star) is an allosteric modulator of nAchR, and its binding to this receptor assures the continuity of the action potential, resulting in muscle contraction and paralysis. Graphic created with BioRender.com. **E**. Levamisole sensitivity assay of wild-type (*n*=129) and FEM-1^Asp133His^ (*n*=122) worms; *N*=3. *n* represents the total number of worms; *N* represents the number of experimental repeats. *p-value* was calculated using the Log-rank (Mantel-Cox) test and is shown adjacent to the graph (ns - not significant). Data was plotted and analyzed in the GraphPad Prism 9 software.

### Abbreviations

Ach: acetylcholine
AchE: acetylcholinesterase
ADHD: attention deficit hyperactivity disorder
ADS: amplicon deep sequencing
ASD: autism spectrum disorder
CCR: constrained coding region
CP: cerebral palsy
CRLs: cullin-RING ligases
DD: developmental delay
E1: ubiquitin-activating enzyme
E2: ubiquitin-conjugating enzyme
E3: ubiquitin ligase
ID: intellectual disability
LOF: loss of function
nAchR: nicotinic acetylcholine receptor
NDDs: neurodevelopmental disorders
NGM: nematode growth medium
MRI: magnetic resonance imaging
UPS: ubiquitin-proteasome system
RING: Really Interesting New Gene
RNA: ribonucleic acid
RNAi: ribonucleic acid interference
RMSD: root-mean-square deviation
WES: whole exome sequencing

