## Supplemental Figure 1 for "A novel *de novo* FEM1C variant is linked to neurodevelopmental disorder with absent speech, pyramidal signs, and limb ataxia"

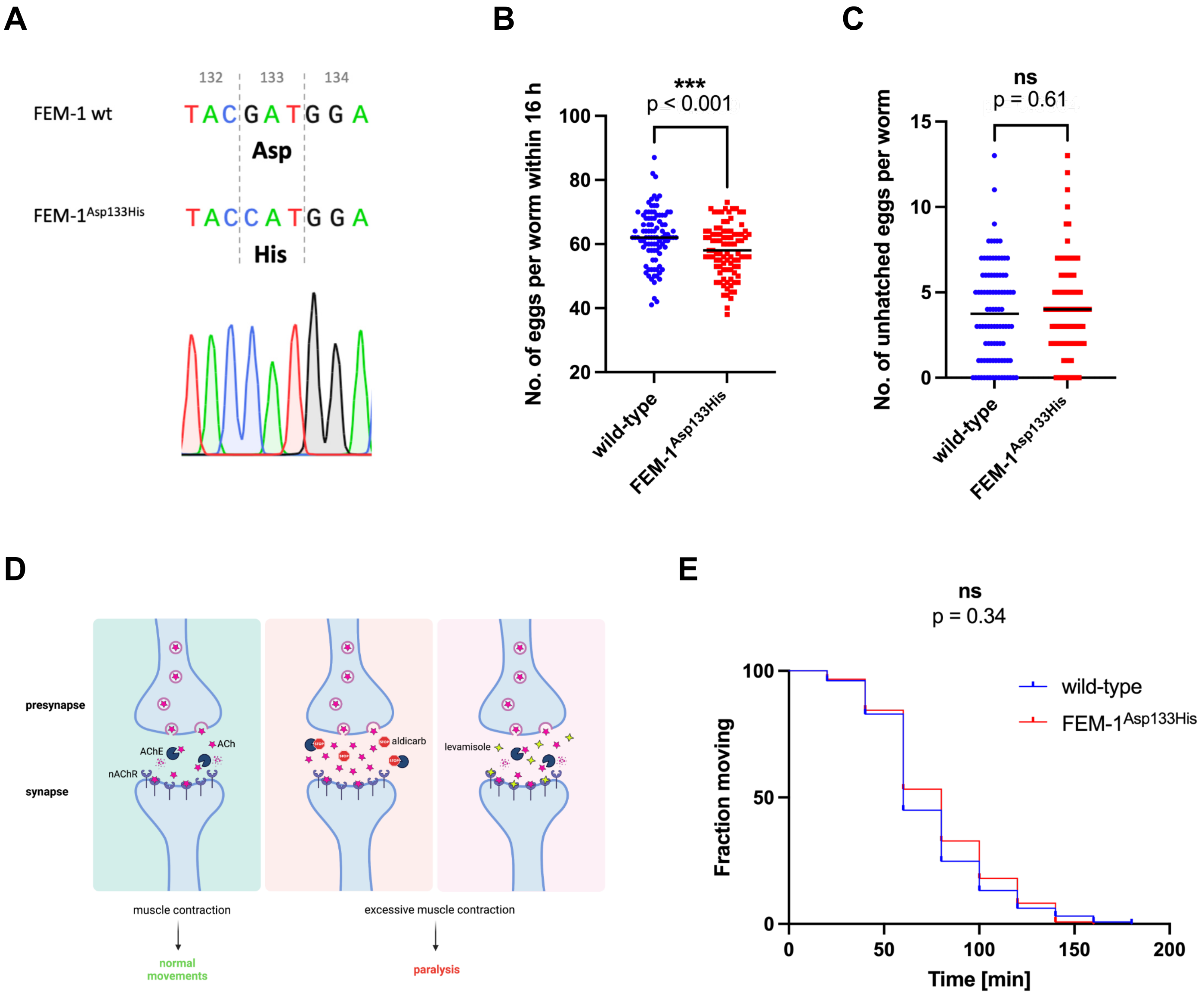


**Supplementary Figure**

**A.** Sequencing chromatogram of CRISPR/Cas9-generated FEM-$1^{Asp133His}$ mutation displaying sequence peaks and base calls. Data visualized in the SnapGene 6 software (Insightful Science; snapgene.com). **B.** Number of unhatched eggs counted after 16 hours of laying for wild-type (*n*=95) and FEM-$1^{Asp133His}$ (*n*=92) worms; *N*=5. *n* represents the number of worms; *N* represents the number of experimental repeats. The stars denote the level of significance of the *p-value* obtained by the Mann–Whitney test. *p-value* is shown adjacent to the graph (***$p \leq 0.001$). Data plotted and analyzed in the GraphPad Prism 9 software. **C.** Egg-laying assay of wild-type (*n=95*) and FEM-$1^{Asp133His}$ (*n*=92) worms; *N=5*. *n* represents the number of worms; *N* represents the number of experimental repeats. *p-value* was calculated obtained by the Mann–Whitney test and is shown adjacent to the graph (ns - not significant). Data plotted and analyzed in the GraphPad Prism 9 software. **D.** Illustration of aldicarb's and levamisole's mechanism of action. The transmission of impulses between neurons is mediated by the synthesis of Ach (magenta star) at the presynapse and its release into the synaptic cleft, where it binds to Ach receptors (only nAchR are shown). Aldicarb (depicted as a stop sign) inhibits AchE (navy pac-man), resulting in build-up of Ach, which causes muscle paralysis. Levamisole (yellow star) is an allosteric modulator of nAchR, and its binding to this receptor assures the continuity of the action potential, resulting in muscle contraction and spastic paralysis. Graphic created with BioRender.com. **E.** Levamisole sensitivity assay of wild-type (*n*=129) and FEM-$1^{Asp133His}$ (*n*=122) worms; *N*=3. *n* represents the number of worms; *N* represents the number of experimental repeats. *p-value* was calculated using the Log-rank (Mantel-Cox) test and is shown adjacent to the graph (ns - not significant). Data plotted and analyzed in the GraphPad Prism 9 software.
